# Statistics of antibody binding to the spike protein explain the dependence of COVID 19 infection risk on antibody concentration and affinity

**DOI:** 10.1101/2021.10.25.465798

**Authors:** David E Williams

## Abstract

The increase of COVID-19 breakthrough infection risk with time since vaccination has a clear relationship to the decrease of antibody concentration with time. The empirically-observed dependence on blood IgG anti-receptor binding domain antibody concentration of SARS-CoV-2 vaccine efficacy against infection has a rational explanation in the statistics of binding of antibody to spike proteins on the virus surface, leading to blocking of binding to the receptor: namely that the probability of infection is the probability that a critical number of the spike proteins protruding from the virus are unblocked. The model is consistent with the observed antibody concentrations required to induce immunity and with the observed dependence of vaccine efficacy on antibody concentration and thus is a useful tool in the development of models to relate, for an individual person, risk of infection given measured antibody concentration. It can be used to relate population breakthrough infection risk to the distribution across the population of antibody concentration, and its variation with time.

## Introduction

The probability of breakthrough infection – where a person who has been previously convalescent or vaccinated becomes infected with Covid19 – is a matter of serious public health concern. The mechanism of antibody neutralisation of viral infection is complex and depends on the type of virus ^1,2^. However, although the variation across individuals is large the concentration in blood of IgG antibodies against the spike receptor binding domain (RBD) of the SARS-CoV-2 virus is well correlated with neutralisation efficacy against the virus ^3^ and appears to be a useful predictor of breakthrough infection risk for both convalescent and vaccinated individuals, regardless of vaccine type ^4-6^. The well-documented increase in breakthrough infection risk over time for some months following vaccination ^7^ has been attributed to a decrease in IgG concentration, in advance of the development later of cell-based immunity ^8^. An empirical model for this dependence has been given^9,10^ and developed into a model describing breakthrough infection risk, and importation risk stratification using quantitative serology ^11^.

Lipsitch et al. ^12^, reviewing recently causes and impact of breakthrough infections in vaccinated individuals, suggest that the rates of breakthrough infection are best viewed as a consequence of the level of immunity at any moment in an individual. This argument can be developed as follows: the probability that an individual becomes infected, *P(infected)*, depends on immune response to previous infection or vaccination, on variant and on contact with other infected people:

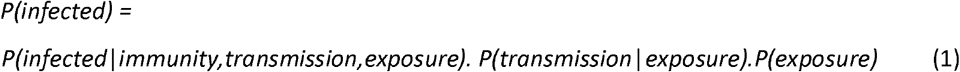

The probability of transmission given exposure, *P(transmission*|*exposure)*, depending on the variant and the nature of contact (close, casual), and *P(exposure)*, depending on the disease prevalence as well as behavioural factors, have been documented through epidemiological studies. Understanding of breakthrough infection thus requires understanding *P(infected*|*immunity,transmission,exposure)*,the probability of infection given all of immunity, transmission and exposure.

Khouri *et al* ^10^ correlated data for vaccine efficacy and antibody concentration across clinical trials for different vaccines. In the presence of antibody, with exposure and transmission probability the same in each group, the vaccine efficacy, *E*, = 1 - (infection proportion amongst vaccinated people / infection proportion amongst unvaccinated people). Khouri et al. used a log-logistic function empirically to derive the dependence of vaccine efficacy against symptomatic infection, *E*, on IgG concentration, *c*, with parameters *c*_*50*_ and k :

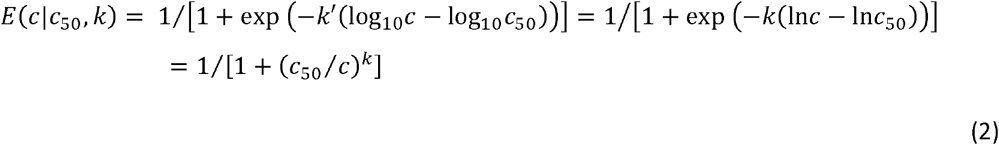

The assumption is that equation (2), derived by correlation of population-median results, also applies at the level of the individual. The risk to an individual at any particular time could then be written:

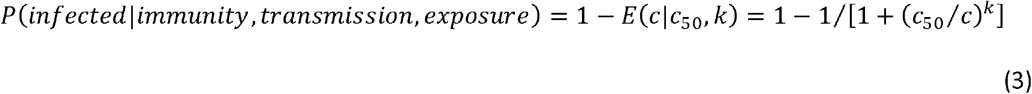

where *c* is the antibody concentration for any given individual at any particular time, *c*_*50*_ depends on the variant and *k* is a parameter independent of variant and vaccine type.

Data on individual infection risk is not available. However, the applicability of equation (3) at the level of the individual can be assessed by using it to understand population breakthrough infection risk. Williams ^11^ used equations (1) and (3) to derive population breakthrough risk given a known antibody concentration distribution across the population, and compared computed and observed population breakthrough infection risk:

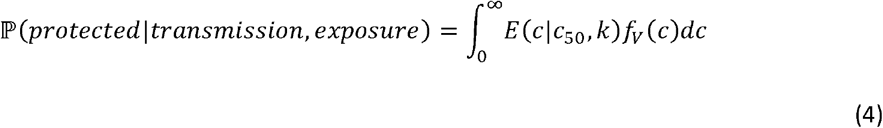

where *f*_*V*_*(c)* denotes the probability density of concentration, *c*, across the population. In a small study, Vargas et al. ^13^indicated that this distribution across a vaccinated population is log-normal, consistent with the observations and assumptions of Khouri et al.^10^

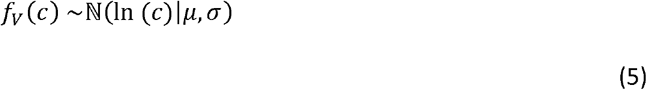

Therefore, if there are sufficient data to estimate the mean, *µ*, and standard deviation, *σ*, over a range of time since vaccination, then an estimate of population vaccine efficacy, *VE*, over this time range can be obtained:

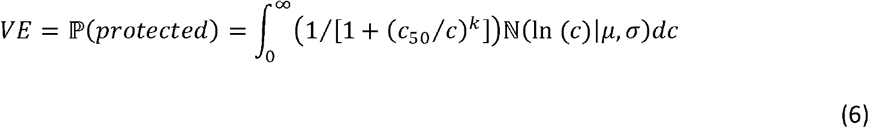

Tartof et al.^7^ in a large retrospective cohort study in California estimated vaccine efficacy for mRNA BNT162b2 COVID-19 vaccine, month-by-month up to 6 months. The study included variants from the original Wuhan strain, against which the vaccine has been derived, up to and including the Delta strain. Israel et al.^14^ measured anti-receptor binding domain (RBD) IgG across a large, BNT162b2 fully-vaccinated population, month-by-month, in Israel. Assuming the concentration distribution is log-normal allows the log-normal mean and standard deviation to be derived from the reported arithmetic mean and variance, median and quartiles. Figure 1 compares the vaccine efficacy reported by Tartof et al.^7^ with the vaccine efficacy computed from equation 6, using the data of Israel et al.^14^, with *c*_*50*_ and k as global fitting parameters averaged over all variants and participant ages (16+). The estimated k = 1.57 (95% CI 1.14, 2.08) agrees with the value deduced by Khouri et al.^10^ Equation (3) appears therefore to be a useful individual risk model with a parameter k that appears to be fixed.

**Figure 1.**
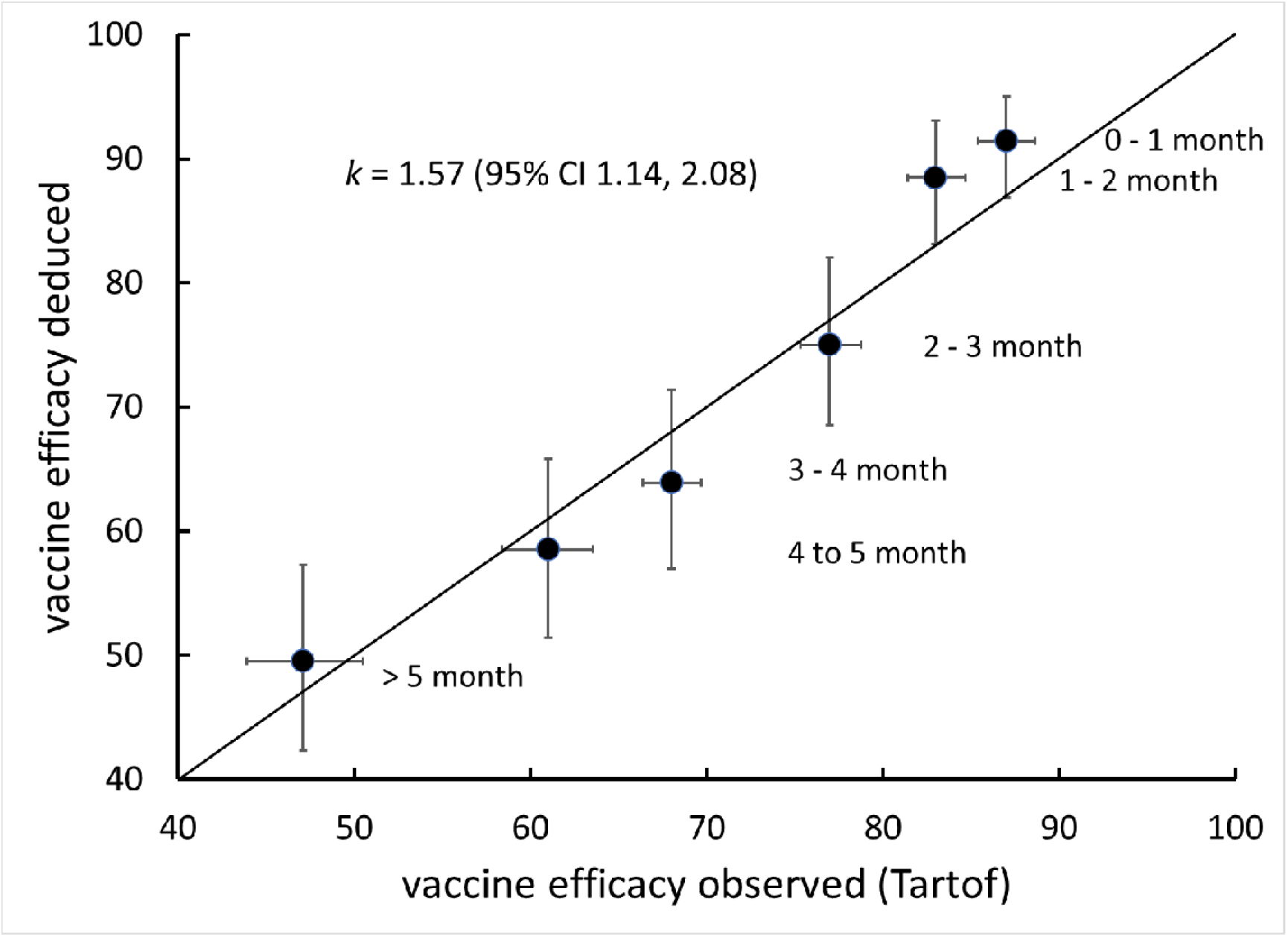
Comparison of vaccine efficacy computed from equation 6 with observed efficacy. Log-normal mean, *µ*, of antibody concentration distributions are the median values given by Israel et al.^14^ Bootstrap sampling from the distribution of *σ* deduced from the various estimates given the data of Israel et al. and from the vaccine efficacy confidence intervals given by Tartof et al.^7^, minimising the sum of squared deviations by varying c_50_ and k. Error bars are 95% confidence intervals.

The model is also useful in assessing the effect of different variants on individual risk. Laboratory and observational studies indicate that, given vaccines directed at the RBD of the Wuhan strain, neutralising efficacy against different variants can be derived simply by scaling the parameter *c*_*50*_ ^9,15-17^. Since the risk model relies heavily on the empirical correlation of vaccine efficacy with neutralising antibody concentration, it would be useful to find a physical basis for the correlation and to use this to develop more confidence in the risk prediction.

## Model

Given current knowledge that vaccine efficacy against infection (as opposed to efficacy against hospitalization and death) is, at least for some months, determined by the concentration of neutralizing antibodies, it is assumed in the following that the mechanism is simply antibody binding to the spike protein blocking the virus binding to host cells ^1,18^. Potent antibodies indeed block binding of the virus to its receptor ^19^.

In the following, we simply suppose that there is some threshold number of spikes on the virus surface that must be unblocked by antibody in order that there is a significant probability that a virus particle may bind to and infect a cell. The total number number, *N*, of spikes per virus particle is variable from one particle to another, distributed over the range 10 – 40 with median around 25 ^20,21^. Let *s* denote the number of antibody molecules bound on a particular particle. We wish to calculate the probability that the number of unoccupied sites is less than or equal to a threshold number, s*: that is that the number of occupied sites is greater than or equal to *N* – *s*\* *P*(*s* ≥ *N*-*s*\*). This probability will depend on the antibody concentration in the medium surrounding the virus particle. The question is therefore : what is the probability distribution for the number of antibody molecules bound per particle as a function of the antibody concentration ?

The problem can be framed in terms of transitions between states of a given particle where each state has a particular number of bound antibodies, ranging from zero up to the maximum, *N*. The objective is to calculate the probability of a given state for a particular particle. The transition frequencies for adsorption and desorption between the different states, *λ*_1,s,_ *λ*_2,s_ depend upon the occupancy, *s* :

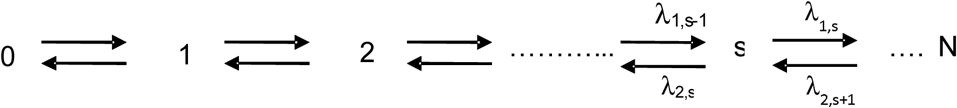

A simplifying assumption is that the diameter of the virus particle and number of spikes/particle are such that the spacing of the spikes is significantly larger than the antibody dimensions so lateral interactions between bound antibodies can reasonably be ignored. With this assumption, the rate of binding of antibody to a particle is proportional to the collision frequency of antibodies with unoccupied sites, hence dependent on the fraction of the particle area that is unoccupied, hence on the fraction of unoccupied sites, whilst the rate of desorption is proportional to the number of occupied sites. Hence for the exchange between state (*s*-1) and state s, where c denotes the solution concentration of antibody

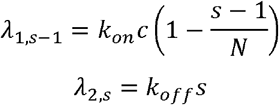

Where *k*_*on*_ and *k*_*off*_ denote the rate constants for attachment and detachment of the antibody to a site on the particle surface. The antibody affinity is the ratio *k*_*on*_ */ k*_*off*_ .

The detail of the calculation is given in the Supporting Information. The key parameter determining the antibody concentration scale for effective blockade is the dimensionless concentration, *z*, which is the product of the antibody solution concentration and the antibody affinity for the binding site. A second parameter depends on *N* and *s\**. The result for the probability that exactly *s* antibody molecules should be bound is:

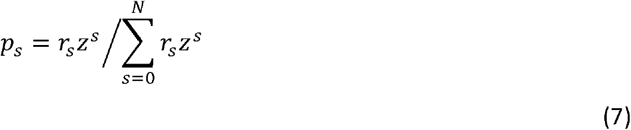

Where

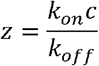

And

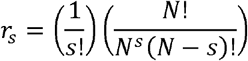

If *N* is significantly greater then *s*, then equation (7) is a Poisson distribution. Therefore, the probability of occupancy *s ≥ N-s\** is:

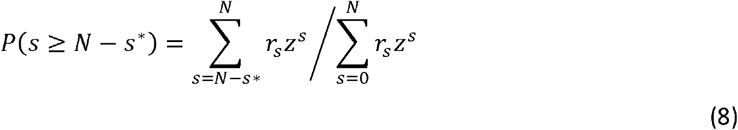

Infection also requires some dose of virus be received. However, as is shown in the following, the dependence of vaccine efficacy on antibody concentration would be just the dependence of *P(s ≥ N-s*)* on concentration, calculated according to equation (8). Infection requires both exposure to an infected person and transmission from that person. Vaccine efficacy is determined by comparing two groups where both the probability of exposure and the probability of transmission given exposure are assumed to be the same. In any given exposure event, the viral dose received, D, would be variable. Then, in the presence of antibody, within some dose, *D*, the number of virus particles that are infectious would be *[1-P(s ≥ N-s*)]D*. Suppose that a ‘critical dose’, *D\**, is required to trigger an infection. Suppose that the dose, *D*, across an exposed population is described by a probability distribution *P(D)*. The probability of infection across the population in the absence of immunity, taking account of the distribution of D would then be 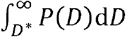In the presence of antibody, the probability of infection would be. 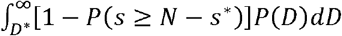 In the presence of antibody, with exposure and transmission probability the same in each group the vaccine efficacy, *E*, = 1 - (infection proportion amongst vaccinated people / infection proportion amongst unvaccinated people). Thus:

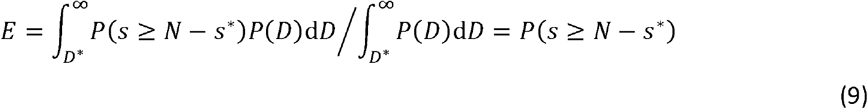

That is: the variation of *E* with antibody concentration, as discussed by Khouri et al.^10^, should be the same as the variation of *P(s ≥ N-s*)* with concentration. Thus we expect

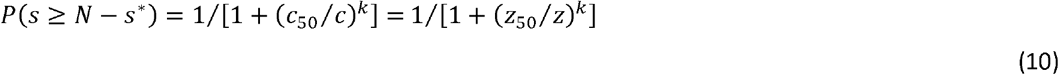

where the dimensionless concentration, *z*, has been substituted.

## Results

Figure 2A shows the variation of *P(s ≥ N-s*)* for various values of *N* and *s\**. The line is fitted to the log-logistic function, demonstrating that the variation of *P(s ≥ N-s*)* with dimensionless concentration, *z*, has the functional form of equation (10), supporting the idea that vaccine efficacy against symptomatic infection can be ascribed simply to antibody blocking of the receptor binding domain of the spike protein. Figure 2C shows that, to obtain high occupancy, *z* must be large – typically z ∼ 1000 - otherwise a significant proportion of particles will have greater than *s\** sites unblocked. Figure 2C also illustrates at the lower values of *z* a potential application of this model to other aspects of the immune response: namely that, if immune recognition requires that a certain minimum number of antibodies be bound to a sufficient proportion of particles then there is a minimum requirement for *z* – typically *z* ≥ 1 .

**Figure 2.**
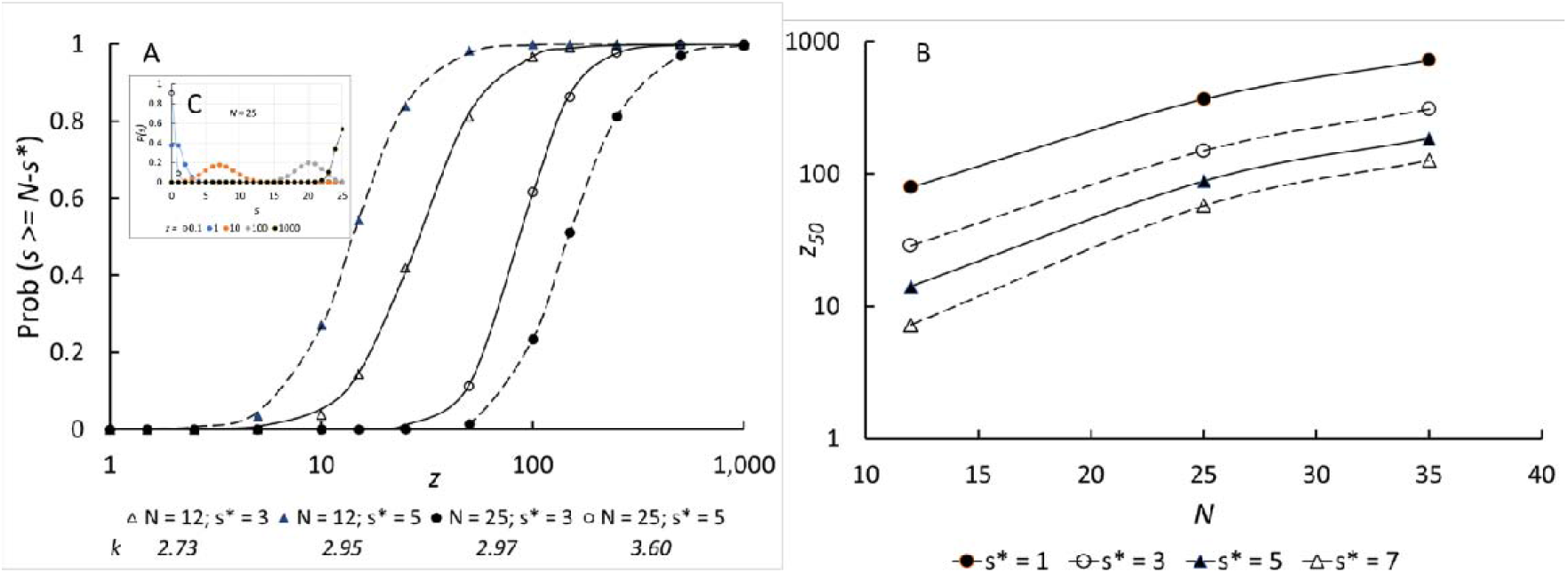
**A**: Probability of site occupancy, *s ≥ N-s*\*, against dimensionless neutralising antibody concentration, *z*, for different numbers of spikes on the virus particle, *N* and various *s*\*; points are calculated and lines are fits to the log-logistic function, equation 10, with rate parameter *k* and dimensionless concentration scaling factor *z*_50_. B: Variation of z_50_ with total binding site number, *N*, and threshold number of vacant sites to allow virus-receptor binding, *s*\*. Inset, C: Individual state probability, *P(s)* against *s*, for different *z*, with *N* = 25

By attributing vaccine efficacy to the probability that more than a critical number of binding sites on the virus should be occupied by antibody, the statistical model captures the observed general behaviour and demonstrates the dependence of the critical parameter, *z*_50_ on the assumption made regarding the critical number of uncovered sites, *s*\*, and on the total number of binding sites / particle, *N*. Since *z* is proportional to antibody affinity, the model captures also the effect of this and attributes the difference between different vaccines, and of the effectiveness of vaccines against different variants, to both the concentration and the affinity of the antibodies induced by vaccination against the receptor binding domain of the different variants. Figure 2B shows that the parameter *z*_50_, interpretable as the median antibody concentration relative to affinity required to achieve 50% blocking, varies strongly both with the number of binding sites, *N*, and the threshold number of unoccupied sites, *s*\*.

Figure 3 shows the variation of *k* determined for the statistical site-binding model for different values of the total number of sites, *N*, that span the range given for the SARS-CoV-2 virus ^20,21^, and with different values assumed for the threshold number of sites left uncovered in order to induce infection, *s*\*. This number is unknown. It may be that virus binding to target requires multiple spike interactions, from spikes that are randomly separated, or may require adjacent spikes, or may be effective with just one spike uncovered. The infection may be ‘land and stick’ or ‘land and seek’ ^22^. The probability that a collision between virus particle and cell is a reactive collision leading to infection would be different for each of these scenarios. In the absence of any evidence to the contrary, given the size scale of the virus the assumption that the spikes are spaced sufficiently far apart that antibody binding to one does not affect binding to another seems reasonable.

**Figure 3.**
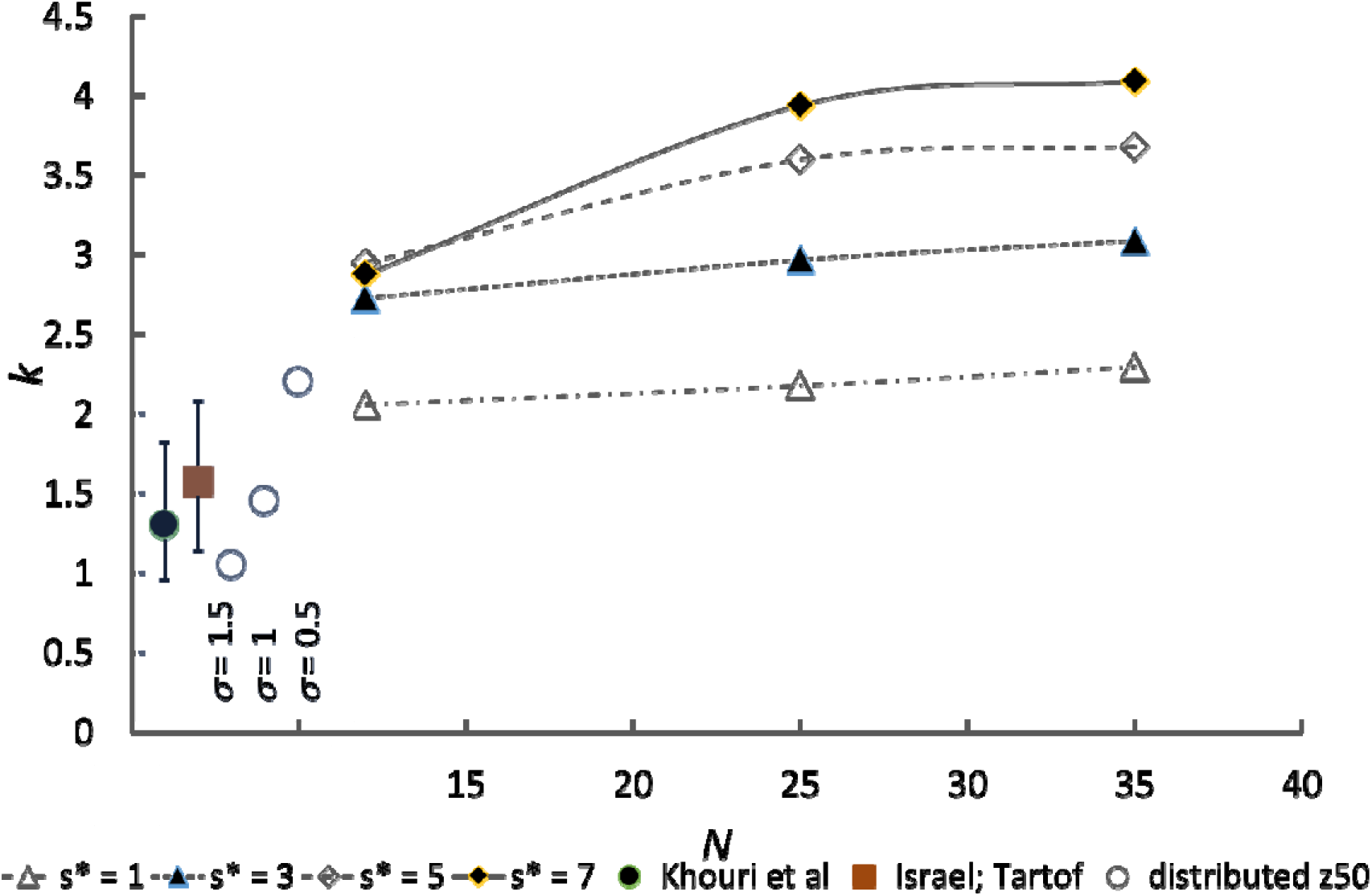
Rate parameter *k* of the log-logistic fit shown in Figure 2 against number of spikes on the virus particle, for different threshold numbers of unoccupied spikes, *s*\*. Symbols ⍰ : effect of introduction of a log-normal distribution of dimensionless concentration, *z*, equivalent to a distribution of neutralising antibody affinity, for spike number *N* = 25 and *N-s*\* = 3; *σ* is the log-normal standard deviation of affinity. ⍰ Comparison with the fit of Khouri *et al*,^10^ describing vaccine efficacy as a function of neutralising antibody concentration, with their 95% confidence interval shown. ⍰ Comparison with the fit of Figure 1 (antibody concentration data of Israel et al.^14^ and vaccine efficacy data of Tartof et al.^7^) with 95% confidence interval from bootstrap sampling.

Khouri et al. give *k* = *k’*/2.303 = 1.30 with 95% confidence interval 0.96 – 1.82, and Figure 1 shows *k* = 1.57 with 95% confidence interval 1.14 – 2.06. The values of the rate parameter, *k*, deduced for different values of *s*\* are rather higher than that deduced by Khouri *et al*, even for the most stringent neutralisation criterion, that only one site unblocked on the virus could lead to infection, although this condition does bring *k* into the range deduced from the data of Tartof et al.^7^ and Israel et al.^14^ (Figure 1 and equation 6). There are two reasons that can be deduced. First, there is a distribution of binding site number. Second, it is known that an antibody population with a range of affinity is induced either by vaccination or by infection ^3,19,23^. The induced affinity distribution may depend on the specific vaccine. Figure 3 shows the effect of variation of *N* on the variation of *k*. For larger *N*, the effect on the variation of k is relatively small. For all *N*, variation of N alone does not bring the computed value of *k* into the range deduced by Khouri et al. or that deduced from Israel et al. and Tartof et al. The effect of a variation of the affinity distribution can straightforwardly be modelled by introducing a distribution of the parameter *z*_50_, whose variation for a particular antibody concentration would be due to variation of antibody affinity. Figure 3 shows as a comparison the effect of introducing a log-normal distribution antibody affinity through a log-normal distribution of *z*_50_. With a distribution that is of moderate broadness, for *s*\* = 3 the deduced value of *k* comes into the middle of the range given by Khouri et al. and to that given on Figure 1, deduced from large studies of breakthrough infection rate and antibody concentration distribution using equation (6). To come to the bottom of the range requires a very broad affinity distribution.

The magnitude scale for antibody concentration can be estimated, as a further qualitative check that the model is sensible. Figure 2 shows that a high degree of protection would require *z*_50_ ∼ 10^2^ – 10^3^. Human antibodies induced in response to SARS-CoV-2 have a range of affinity (ratio of ‘on’ rate constant to ‘off’ rate constant, *k*_*on*_*/k*_*off*_) with the most potent ∼10^11^ M ^19^, to the receptor binding domain. Thus, given the deduced range of *z*_50_, the expected range of median antibody concentration would be ∼10^−9^ – 10^−8^ M. Data from Roche ^24^ indicate median convalescent antibody concentration ∼ 4 nM and from Wei *et al*^25^. post-vaccination concentrations in the range 200 – 500 ng / mL (1.5 – 3.5 nM assuming an antibody molecular weight of 150 kDa) whilst other studies (converting units) show concentrations above 10 nM. The antibody concentration range deduced from the model therefore seems reasonable.

## Conclusion

The empirically-observed dependence of vaccine efficacy on antibody concentration^10^ has a rational explanation in the statistics of binding of antibody to spike proteins on the virus surface. The model is consistent with the observed antibody concentrations required to induce immunity and with the observed dependence of vaccine efficacy on antibody concentration. It provides a way to constrain the value of the parameter describing the increase of vaccine efficacy with increase of antibody concentration and thus is a useful tool in the development of models to relate, for an individual person, risk of breakthrough infection given measured antibody concentration. It provides an explicit means for relating laboratory measurements of the antibody affinity for virus neutralisation or spike protein binding to expected breakthrough infection rates and hence should be useful in understanding the public health implications of new variants.

## Supporting information

derivation of state probabilities

