## Supplementary material for "Statistics of antibody binding to the spike protein explain the dependence of COVID 19 infection risk on antibody concentration and affinity": derivation of state probabilities

**Supporting Information**

### COVID 19 breakthrough infection risk: a simple physical model describing the dependence on antibody concentration

David E Williams

School of Chemical Sciences, University of Auckland, Private Bag 92019, Auckland, 1142, New Zealand

#### Calculation of the occupancy probability

The occupancy probability, *p(s,t)* , is determined as follows [1]:

[probability that occupancy of a particle is *s* at time *t* ] =

[probability that the occupancy was *(s-1)* at *(t-δt)* and that one further antibody was captured in the interval *(t-δt)* to *t* ]

+ [probability that the occupancy was *(s+1)* at *(t-δt)* and that one antibody was desorbed from the surface in the interval *(t-δt)* to *t* ]

— [probability that the occupancy was at *s* at *(t-δt)* and that antibody was captured or lost in the interval *(t-δt)* to *t* ]:

$$p(s,t) = p(s-1,t-\delta t)\lambda_{1,s-1}\delta t + p(s+1,t-\delta t)\lambda_{2,s+1}\delta t + p(s,t-\delta t)(1-\lambda_{1,s}\delta t - \lambda_{2,s}\delta t)$$

Hence, altering the notation : *p(s,t)* = $p_s$ , and substituting for $\lambda$

$$\frac{dp_s}{dt} = p_{s-1}\lambda_{1,s-1} + p_{s+1}\lambda_{2,s+1} - p_s(\lambda_{1,s} + \lambda_{2,s})$$

$$= p_{s-1}k_{on}c\left(1-\frac{s-1}{N}\right) + p_{s+1}k_{off}(s+1) - p_s\left(k_{on}c\left[1-\frac{s-1}{N}\right] + k_{off}s\right)$$

\- (1)

For the state 0, from which there is no antibody desorption,

$$\frac{dp_0}{dt} = p_1k_{off} - p_0k_{on}c$$

(2)

And for the state *N*, from which there is no further antibody adsorption,

$$\frac{dp_N}{dt} = p_{N-1}k_{on}c\left(1-\frac{N-1}{N}\right) - p_Nk_{off}$$

(3)

The initial condition is: $p(0) = 1$ at $t = 0$ and furthermore $\sum_{s=0}^{N} p_s = 1$ at any $t$.

The solution for the time-varying probabilities can be obtained numerically. The solution for the steady-state occupation probability is obtained by setting the derivatives to zero and solving recursively starting with the determination of $p_1$ from eq(2), applying eq (1) to obtain successively the $p_s$, and applying $\sum_{s=0}^{N} p_s = 1$ to obtain $p_0$.

Thus, defining $z = \frac{k_{on}c}{k_{off}}$ and $r_s = \left(\frac{1}{s!}\right)\left(\frac{N!}{N^s(N-s)!}\right)$ gives:

$$p_s = r_s z^s / \sum_{s=0}^{N} r_s z^s \quad (3)$$

If $N$ is very large and the total occupancy is sufficiently small, then $p_s$ follows a Poisson distribution: $p_s = \frac{z^s}{s!} exp(-z)$.


1. Ishida, K., Stochastic Model for Langmuir Isotherm. *Bulletin of the Chemical Society of Japan* **1969,** *42* (2), 563-564.
